# FastField: An Open-Source Toolbox for Efficient Approximation of Deep Brain Stimulation Electric Fields

**DOI:** 10.1101/2020.03.03.974642

**Authors:** Mehri Baniasadi, Daniele Proverbio, Jorge Gonçalves, Frank Hertel, Andreas Husch

## Abstract

Deep brain stimulation (DBS) is a surgical therapy to alleviate symptoms of certain brain disorders by electrically modulating neural tissues. Computational models predicting electric fields and volumes of tissue activated are key for efficient parameter tuning and network analysis. Currently, we lack efficient and flexible software implementations supporting complex electrode geometries and stimulation settings. Available tools are either too slow (e.g. finite element method–FEM), or too simple, with limited applicability to basic use-cases. This paper introduces FastField, an efficient open-source toolbox for DBS electric field and VTA approximations. It computes scalable e-field approximations based on the principle of superposition, and VTA activation models from pulse width and axon diameter. In benchmarks and case studies, FastField is solved in about 0.2s, ~ 1000 times faster than using FEM. Moreover, it is almost as accurate as using FEM: average Dice overlap of 92%, which is around typical noise levels found in clinical data. Hence, FastField has the potential to foster efficient optimization studies and to support clinical applications.

## 1. Introduction

Deep brain stimulation (DBS) is a neurosurgical method to electrically stimulate specific brain regions. It is an established therapy for Parkinson’s Disease, Essential Tremor and Dystonia (Deuschl et al., 2006; Flora et al., 2010; Larson, 2014) and is emerging for several other diseases like Obsessive-Compulsive Disorder (Abelson et al., 2005) and Anorexia nervosa (Wu et al., 2013). The procedure is based on implanting electrodes (or “leads”) delivering electrical pulses to the neural tissue. There are several lead designs available, providing a recently increasing complexity of possible contact arrangements, including segmented leads (Buhlmann et al., 2011; Horn et al., 2019). Some of the current widely-used electrode geometries are shown in Fig. 2. Augmented complexity allows for better targeting of disease-specific brain regions (FDA, 2015; Lee et al., 2019), while avoiding areas associated with side effects (Mallet et al., 2007).

Simulating the propagation of induced electric fields (e-field) enables prediction of the DBS effects on neural tissue (Anderson et al., 2018; Åström et al., 2015; Butson and McIntyre, 2008; Cubo, 2018; Horn et al., 2017, 2019; McIntyre and Grill, 2002). The portion of tissue affected by a propagating e-field is typically quantified by the “volume of tissue activated” (VTA). VTA is a conceptual volume that is thought to elicit additional action potentials due to electrical stimulation of axons (McIntyre and Grill, 2002). It is usually identified by a threshold value *T* to define iso-surfaces of effective e-field (Åström et al., 2015).

### 1.1. Limitations of current DBS simulations

Reconstructing electric fields in the brain is complex, primarily due to its heterogeneity. Apart from skull and skin, the brain is mostly composed by white matter (WM), grey matter (GM) and cerebrospinal fluid (CSF), which features different tissue properties like electrical conductivity (Howell and McIntyre, 2016). White matter in particular, having a considerable amount of fibre tracts, influences the spatial propagation of electric fields (Gabriel et al., 2009; Suh et al., 2012). To improve model accuracy, information about patient-specific white matter anisotropy can be extracted from diffusion tensor images (DTI) (Butson et al., 2007). Additionally, models may include dielectric dispersion and other details of the medium.

Currently, the most flexible and detailed computational models, that also consider complex electrode designs, are based on Finite Element Methods (FEM) (Åström et al., 2015; Cubo, 2018; Horn et al., 2017; Howell and McIntyre, 2016). They partition the brain into finite sets of basic elements (typically tetrahedrons), each potentially parametrised with tissue-specific conductivity values. However, despite the vast literature, there is still no global consensus on conductivity values of certain brain tissue classes (cf. Table 1 and references therein).

**Table 1.** Conductivity values [S/m] for different tissues reported in the literature. **Left:** heterogeneous medium, with values for white matter (WM), grey matter (GM) and Cerebrospinal Fluid (CSF). Values refer to the most recent literature. The spanned interval is considerable: values range from 0.058 S/m to 0.14 S/m for white matter, 0.089 S/m to 0.33 S/m for grey matter, and 1.5 S/m to 2 S/m for CSF. **Right:** conductivity values [S/m] when the brain is treated as a homogeneous medium. They range from 0.1 S/m to 0.2 S/m. Values refer to the most recent literature.

| Heterogeneous medium |  |  |  | Homogeneous medium |  |
| --- | --- | --- | --- | --- | --- |
| WM | GM | CSF | Reference | Values | Reference |
| 0.058 | 0.089 | 2 | (Cubo et al., 2019) | 0.1 | (Åström et al., 2015) |
| 0.059 | 0.0915 | - | (Horn et al., 2019) | 0.1 | (Cubo and Medvedev, 2018) |
| 0.06 | 0.15 | 1.79 | (Cendejas Zaragoza et al., 2013) | 0.123 | (Alonso et al., 2018) |
| 0.075 | 0.123 | 2 | (Alonso et al., 2018) | 0.2 | (Howell and Grill, 2014) |
| 0.075 | 0.123 | 2 | (Hemm et al., 2016) | 0.2 | (Vorwerk et al., 2019) |
| 0.14 | 0.23 | 1.5 | (Howell and McIntyre, 2016) | 0.2 | (Anderson et al., 2018) |
| 0.14 | 0.33 | - | (Horn et al., 2017) |  |  |
| 0.14 | 0.33 | 1.79 | (Vorwerk et al., 2019) |  |  |

Overall, complex FEM-based models (Butson and McIntyre, 2008) are powerful at estimating DBS electric fields and VTA, but they suffer from high computational costs. This slows down multiple parameters testing and hinders computational optimization (Cubo et al., 2019). It also limits clinical application, as physicians require rapid responses. Moreover, their precision is often shadowed by noise and finite precision of real measurements.

To simplify DBS reconstructions, several tools approximate the brain as a homogeneous medium (Alonso et al., 2018; Anderson et al., 2018; Åström et al., 2015; Cubo and Medvedev, 2018; Howell and Grill, 2014; Vorwerk et al., 2019). Table 1 (right) contains commonly used conductivity values. Other simplifications include fully heuristic models that directly estimate VTA shapes from stimulation parameters, without explicitely simulating the electric field (Chaturvedi et al., 2013; Dembek et al., 2017; Kuncel et al., 2008; Mädler and Coenen, 2012). These models are fast, but they only support ring-shaped contact designs and mono-polar stimulation.

### 1.2. FastField

The aim of this work is to introduce a flexible and efficient algorithm addressing the drawbacks of currently available software. Indeed, FastField estimates DBS induced electric fields in the order of milliseconds. It supports complex electrode designs and is easily extendable for future geometries. It also provides an activation model for VTA considering different pulse widths and axon diameters, while preserving the quick timing. FastField predictions are nearly as accurate as FEM-based models with homogeneous conductivity for the brain and different conductivity values for conducting and isolating parts of the electrode. Its main contribution is thus being a comprehensive trade-off between accuracy of simulations and rapid response. It is provided as an open-source toolbox and the graphical user interface and the source code are freely available for public use. Hence, FastField is applicable in clinical practice (to test different configurations) and in optimization studies. Its computational workflow is presented in Fig. 1.

**Fig. 1.**
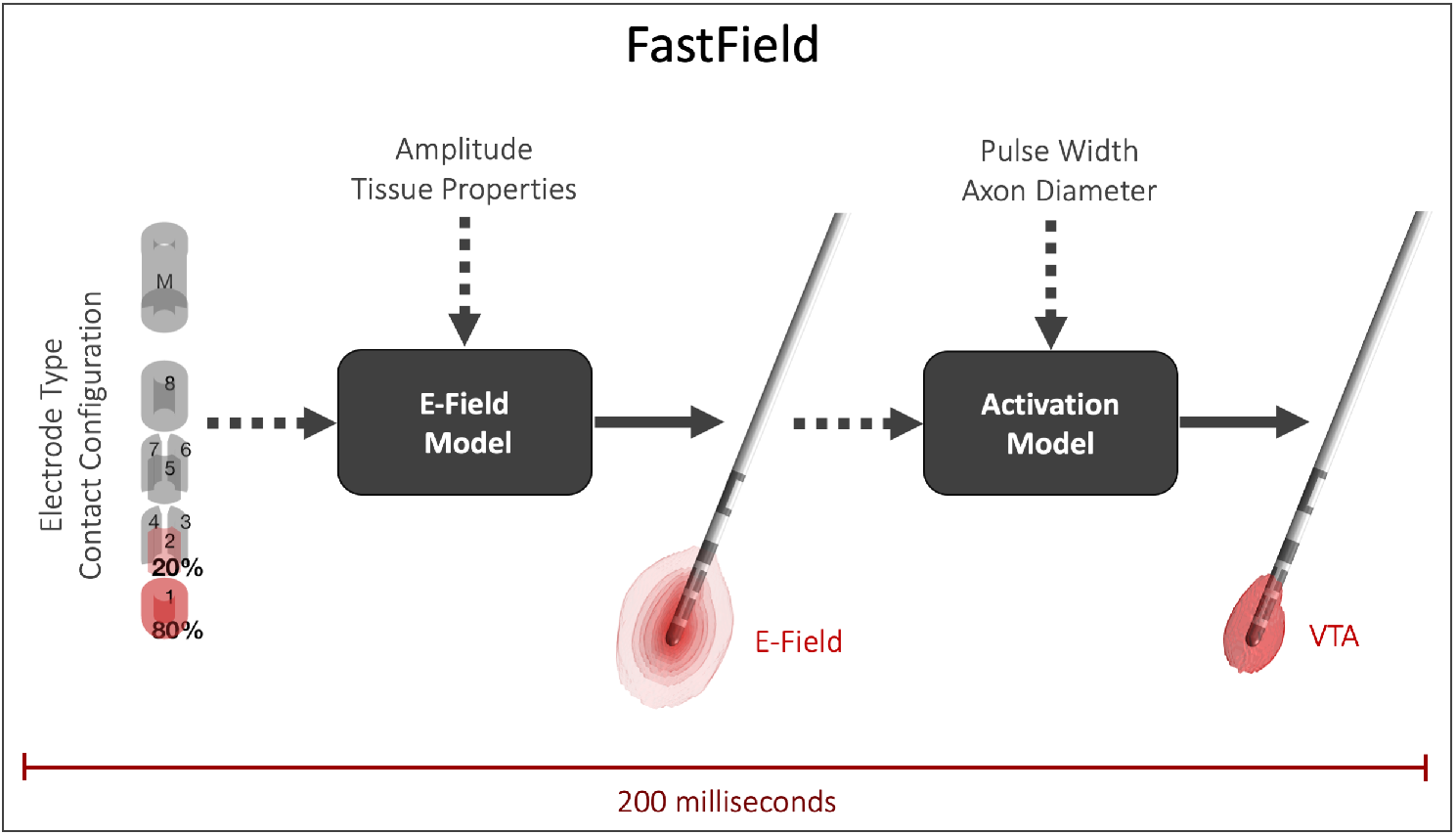
FastField workflow. FastField consists of two independent stages: a fast e-field estimation followed by a heuristic prediction of the VTA. Inputs for the e-field model are the electrode contact configuration, stimulation parameters and assumed tissue properties. Patient’s electrode location in MNI space may be added for patient-specific studies. The subsequent VTA estimation allows to consider different pulse widths and axon diameters. The whole process is fully automatic and takes about 0.2 s on a standard computer.

## 2. Methods

FastField inputs are: electrical conductivity [S/m], the stimulation amplitude ([mA] or [V] depending on the machine setting) and contact configuration, i.e. the active contacts and their relative weight. FastField then calculates the strength of the electric field on a standard grid around the electrode (Sec. 2.2) from inputs and a group of pre-computed e-fields (cf. Sec. 2.1). To estimate the e-field threshold for the VTA, FastField activation function also considers the stimulation pulse width and the hypothesised axon diameter (Sec. 2.3).

To personalise the simulation, the patient’s electrode location in MNI space in Lead-DBS format can be added (more in Sec. 2.4). Target structures are extracted from a brain atlas registered into the MNI space for final visualization (Sec. 2.5). The toolbox has a user-friendly GUI for practical use (Sec. 2.6).

Finally, we introduce two metrics to gauge the accuracy resulting from e-filed approximation (Sec. 2.7).

### 2.1. Standard e-field library

Standard e-field library (or “pre-computed e-fields”) is derived from finite element models where only one contact of the electrode is active at a time, for different geometries (Fig. 3). First, a cylinder domain is defined around the electrode. The area inside the cylinder is divided into three regions: brain, conducting and insulating part of the electrode. Tetrahedron meshes are generated and linked to regions where different electrical conductivity is assumed (brain area: *κ* = 0.1 S/m; conducting electrode parts: *κ* = 10^8^ S/m; insulating electrode parts: *κ* = 10^−16^ S/m). The electric field strength [V/mm] is simulated at the center of each mesh for constant current *A*_0_ = 1 mA. This procedure is repeated for each contact of all electrode types (cf. Fig. 2).

**Fig. 2.**
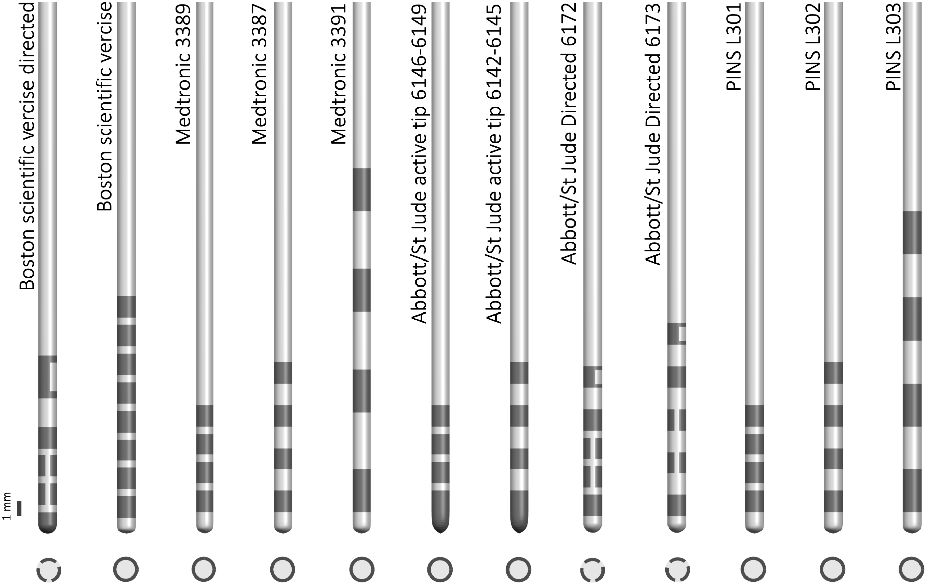
Common DBS electrode geometries. Lateral (top) and longitudinal (bottom) views of the electrodes are shown. Medtronic 3389, 3387, and 3391 (Medtronic, Dublin, Ireland), St Jude Medical (Abbott Laboratories, Abbott Park, Illinois, USA) active tip 6146-6149 and 6142-6145, and PINS Medical L301, L302, and L303 (Beijing, China) have 4 rings of conductive contacts; Boston Scientific (Marlborough, Massachusetts, USA) vercise has 8 rings. Boston scientific vercise directed, St Jude Medical Infinity Directed 6172 and 6173 have 2 full rings and 2 rings segmented into 3 conductive contacts. Note that the size and the distance between the contacts also differ between the leads (Okun et al., 2012; Schuepbach et al., 2013; Timmermann et al., 2015)

**Fig. 3.**
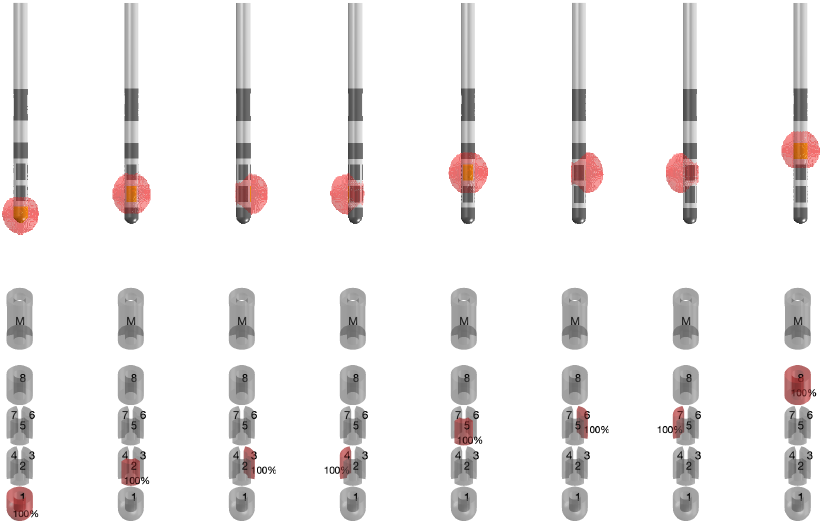
Standard e-field library for Boston scientific vercise directed for constant-current. On top, the simulated e-fields with Simbio/FieldTrip FEM model are shown; below, the corresponding contact configurations. This electrode has 8 conductive contacts, so 8 e-fields are simulated (one for each contact). Default amplitude is *A*_0_ = 1 mA. Similarly, standard e-field libraries for other electrode types are generated.

The above preliminary computations are performed with Lead-DBS Simbio/FieldTrip (Horn et al., 2019). Next, Lead-DBS interpolating function converts the e-field values from the arbitrary mesh to a 3D grid of constantly spaced points. The grid 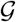 is referred to as “standard grid” and is used as a common template. By convention, 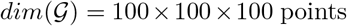 (average point distance is 0.2 in [mm]). Pre-computed e-field values on 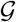 are finally stored in the standard e-field library.

Real devices allow voltage [V] as input setting. Hence, the algorithm allows conversion to amplitude units, considering the device impedence as additional input.

### 2.2. FastField computation

FastField algorithm simulates the electric field on the standard grid. For each contact, the corresponding library is initially chosen based on the amplitude mode and the electrode type. Then, FastField scales the pre-computed e-field by the weighted activation amplitude of the corresponding contact and by the user-defined brain conductivity. Finally, it computes the total e-field *E*(*g*) by exploiting the additive property of electric fields (in line with Anderson et al. (2018); Slopsema et al. (2018)). Formally, *E*(*g*) is computed at each point *g* of the 3D grid 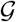 as:

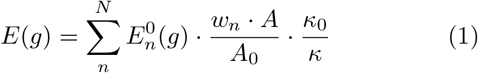

Here, *N* is the number of contacts of the electrode, subscript *n* identifies each contact. 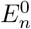 is the pre-computed e-field for each contact with weight *w_n_*. *A* and *κ* are amplitude and conductivity defined by the user, *A*_0_ and *κ*_0_ are amplitude and conductivity used to generate the standard library and are equal to 1 mA and 0.1 S/m.

To smooth the electric field on the grid, convolution is performed with a Gaussian kernel. Next, a system of linear equations is solved for the 4 marker coordinates (head, tail, X, and Y, cf. 2.4) to get the transformation matrix *M* to MNI space. The standard grid is thus transformed and tilted with respect to the position of the patient’s electrode, that is placed at the center of the transformed grid. Finally, the target location is extracted from the combined atlas (Sec. 2.5) for the final visualization.

### 2.3. A flexible model for the Volume of tissue activated

Current open-source models only provide a small set of parameter combinations to compute the stimulation field threshold *T* for the volume of tissue activated. In FastField, we implement a straightforward heuristic model to fit published data on pulse width *P_W_*, axon diameter *D* and resulting e-field threshold *T* (*P_W_*, *D*). The latter defines the iso-surface of the VTA.

The model is obtained as follows. We first develop a heuristic simplification of the axon electrical and geometrical properties. Considering a heterogeneous manifold of axons in the region around the DBS lead, our minimal model refers to the mean properties of such a manifold and not to the particular geometry or conductivity of a single axon. Hence, instead of considering complex geometries as in Åström et al. (2015), we approximate a “mean field” axon with a cylindrical conducting cable. In addition, we consider the conductance along the cable as closely ruled by Ohm’s law. In this sense, *V_T_* (*P_W_*, *D*) is the electric potential along the cable. Then *E_T_* = Δ*V_T_* is its gradient, commonly referred to as the electric field strength. In turn, *T* (*P_W_*, *D*) approximates the threshold for axonal activation under the effect of *E_T_*. It is proportional to the product of *P_W_* (providing energy, cf. Dembek et al. (2017)) and *D*, that influences the conductance and thus the dampening of electric signal. Because of heterogeneity in shape and electrical properties, the functional dependence is scaled by power laws to be fitted with available data. The heuristic model reads:

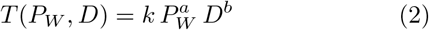

To enable a straightforward fit in the Matlab Curve Fitting toolbox, we then convert the log-linear fit for the model into an exponential form (*c* = log *k*):

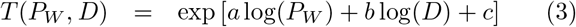

The FastField algorithm thus allows the user to define the desired threshold value with extended flexibility, that is, also considering pulse width and axon diameter. Thanks to the heuristic model, the quick timing is preserved. Calibration of the model with published data and subsequent *in silico* experiments are reported in Sec. 3.1.

### 2.4. Patient’s pre-processing

Evaluating patient’s data requires the electrode position in MNI space. Thus, we perform the following preprocessing steps. Patient’s Computed Tomography (CT) scan and T1- and T2-weighted Magnetic Resonance Imaging (MRI) are linearly registered to each other and nonlinearly to MNI space. We use Advanced Normalization Tool (ANTs, http://stnava.github.io/ANTs/) and FMRIB’s Linear Image Registration Tool (FLIRT) (Ashburner, 2007; Avants et al., 2008; Jenkinson et al., 2002; Jenkinson and Smith, 2001) for patient’s MRI and CT scan registration, respectively. Then, the PaCER algorithm (Husch et al., 2018) returns the location of the electrode in the brain, while the DiODe algorithm returns its rotation (Hellerbach et al., 2018). By this combination, we estimate the head, tail, X and Y coordinates of the marker (reference label on the lead). With these, we calculate the transformation matrix from the standard electrode space into MNI space considering the patient’s electrode location.

### 2.5. Combined atlas

There are several brain atlases registered into MNI space. Distal atlas is explicitly generated for Lead-DBS use (Ewert et al., 2018). However, distal atlas does not contain all DBS target structures, e.g. nucleus accumbens that is included in CIT168 atlas (Pauli et al., 2018). Therefore, the FastField build-in library combines both Distal and CIT168 atlas.

### 2.6. The graphical user interface

FastField graphical user interface is shown in figure 5. It is designed so to provide a comfortable user experience. Input settings are located on the left-hand side of the GUI, while the output location of the electrode in the brain and the VTA are shown on the right-hand side. Additional options for visualization are also present.

**Fig. 4.**
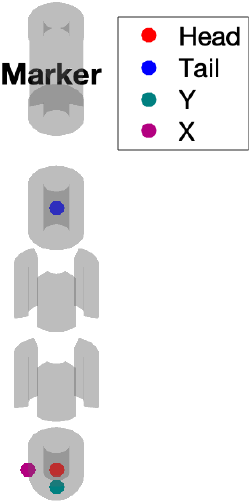
Head, tail, X, Y marker coordinates on Boston scientific vercise directed lead model. These points are used to locate the electrode in MNI pace. Conventionally, head is the center point of the lowermost contact, and tail the center point of the uppermost contact. To locate X and Y, consider a plane perpendicular to the electrode shaft, passing by the head point. The point on the plane that has the least distance to the center of the marker is the Y point. X is perpendicular to the line passing by head and Y.

**Fig. 5.**
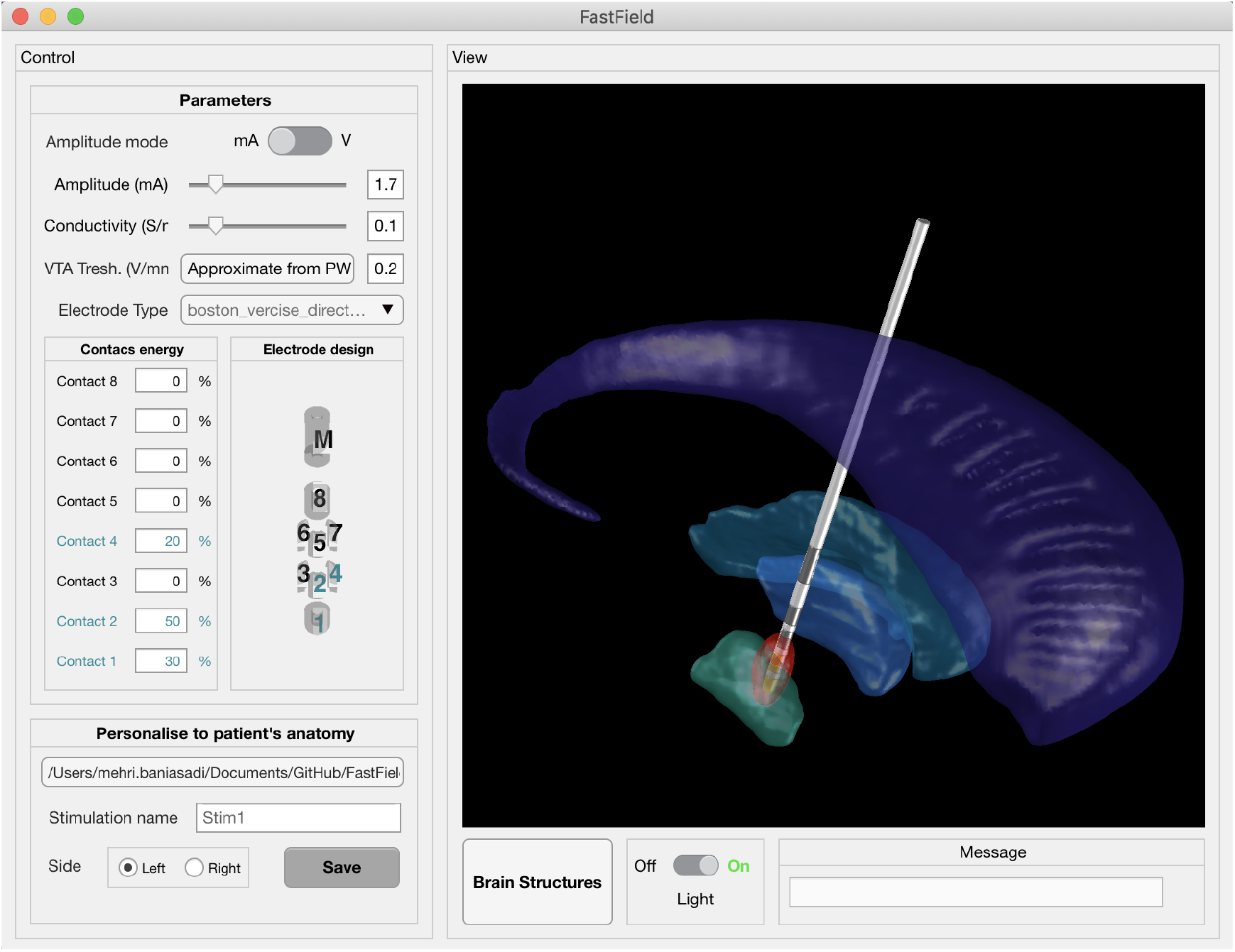
FastField graphical user interface. Left panel includes input values, VTA threshold estimation button and contact configuration for the chosen lead. Right panel is for output visualization. Additional panels allow navigation of patient’s data and additional settings for visualization. As an example, input values are set as follows: amplitude *A* = 1.7 mA, conductivity *κ* = 0.1 S/m, threshold *T* = 0.2 V/mm, pulse width *P_W_* = 60*μ*s and axon diameter *D* = 3.4*μ*m. The electrode type is Boston scientific vercise directed. 30 % of the energy is on contact 1, 50% on contact 2 and 20% on contact 4. STN, internal globus pallidus(GPi), external globus pallidus(GPe) and Caudate are the visualized structures in light green, blue, dark green, and purple. The VTA, here from a general heuristic value *T* = 0.2 V/mm (as suggested by Horn et al. (2017) based on Hemm et al. (2005)) is shown in red.

Main inputs are: stimulation amplitude, brain tissue conductivity, type of electrode, contact configuration and the percentage of energy on each contact. Stimulation amplitude can be set in [mA] or in [V] according to the machine settings. Additionally, VTA threshold can be estimated in a pop-up window by specifying pulse width and axon diameter. These inputs can be directly used in abstract studies that estimate the general effects of different electrodes and contact configurations without being patient-specific.

For patient-specific studies, users may provide a dedicated folder containing the patient’s electrode location in MNI space. The corresponding file should include the position of the electrode marker, including 4 points of head, tail, X and Y (Sec. 2.4). The user can then visualize the electric field by changing the main inputs as described above. Different brain regions can also be visualized, to evaluate the structures affected by the e-field. Finally, the electric field information can be easily exported for further studies.

### 2.7. Accuracy measurement

FastField relies on an approximated estimation of the electric field within the brain. It is then informative to quantify how it differs from more complete finite element models. We do so by computing the absolute deviation between our e-field (*E*_1_) and a reference e-field (*E*_2_), for each point *g* of the same template grid 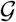. The sum of the absolute deviation values over 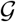 is then normalized on the global strength of the reference field, thus estimating the relative error:

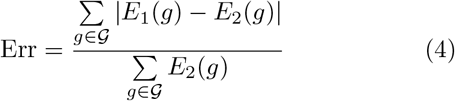

We then call “accuracy” of the FastField simulation, with respect to reference FEM-based field, the quantity:

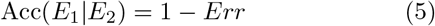

Several experiments with different electrode types and settings are reported in Sec. 3.2.

To estimate the similarity between FastField and FEM predictions, we also compute the Dice score metric on two VTAs (A and B), defined as:

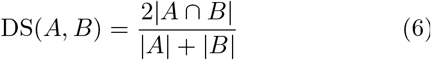

where |*A*| and |*B*| are the cardinalities of the two sets.

## 3. Results

The VTA model calibration is presented in Sec. 3.1. Next, the results of FastField are benchmarked against a realistic FEM-based model to estimate the accuracy (cf. Sec. 3.2). We also present three case studies to illustrate the practical application of our algorithm (cf. Sec. 3.3, 3.4,3.5). Details on data acquisition and management are commented at the end of the paper.

### 3.1. Calibrating the VTA model

The volume of tissue activated model (Eq. 3) is fitted to data published in Table 3 of Åström et al. (2015) in a non-linear least-squares sense using Matlab Curve Fitting Toolbox. These data are reported to be accurate for a stimulation voltage of 3*V*.

Figure 6 visualizes the fitted model surface for pulse widths *P_W_* ∈ [1; 240] *μS* and axon diameters *D* ∈ [1; 8]*μm*. Calibrated values for the model coefficients *a, b, c* are also reported in the figure. The goodness of fit is estimated by considering a reduced R-square statistics over the degrees of freedom. In this case, *R*^2^_*reduced*_ = 0.9948 ~ 1. Both the general heristics of *T* = 0.2*V/mm* and an additional experimental point (Åström et al., 2015) lie within the surface, thus strengthening its validity for practical use. Direct use of the developed heuristic model to estimate the isocontour lines for the volume of tissue activated is shown in Fig. 7. In there, comparison with a full electric field computed by FEM model SimBio/FieldTrip is also reported. The heuristic model increases FastField flexibility by considering various *P_W_* and *D*, without increasing its computational load. This aspect also allows for direct comparison of different settings, thus extending the testable parameters and the application of the algorithm in abstract studies and clinical practice.

**Fig. 6.**
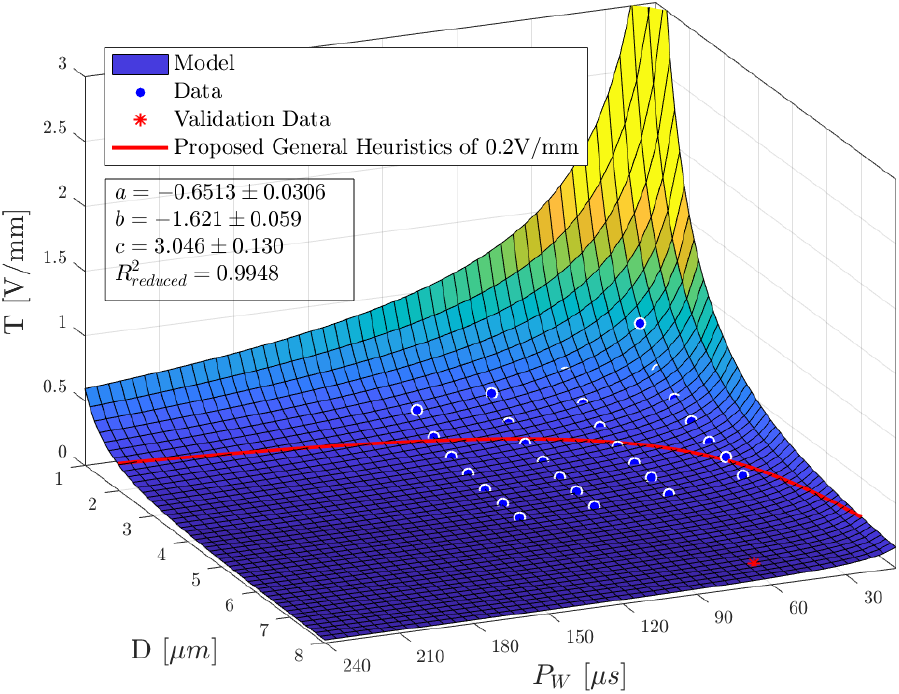
**Plot of the VTA model surface**, predicting the threshold *T* given pulse width *P_W_* and axon diameter *D*. Data from Table 3 in (Åström et al., 2015) used for fitting are visualized as circles. An additional point reported in Table 2 in (Åström et al., 2015) used for validation is denoted by an asterisk. The isocontour of the common general heuristics of *T* = 0.2V/mm as suggested by Horn et al. (2017) based on (Hemm et al., 2005) is denoted in red. Calibrated parameters and goodness of fit are listed in the textbox.

**Fig. 7.**
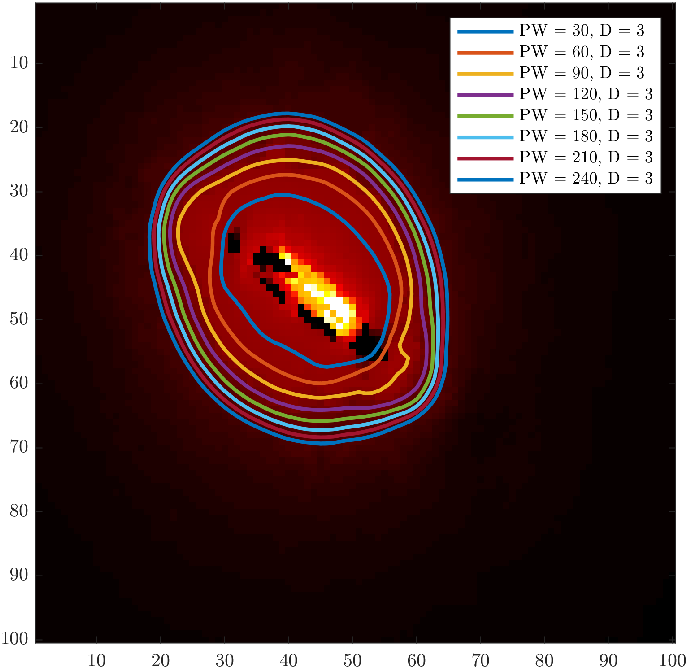
**Overlay of e-field threshold isocontour lines** as predicted by our model for different values of pulse width at a constant axon diameter. On the background (red area), e-field of a Boston Scientific electrode simulated using SimBio/FieldTrip as implemented in Lead-DBS.

### 3.2. FastField Accuracy

We compare the electric field estimated with FastField with the one simulated with Lead-DBS Simbio/FieldTrip finite element model, on the same template domain. We consider different electrode types and DBS settings, including different contact configurations and amplitude values. For simulations with Simbio/FieldTrip method, there are two scenarios for *E*_2_: heterogeneous medium with Lead-DBS default conductivity values (*κ* = 0.132 S/m for grey matter and *κ* = 0.08 S/m for white matter) and homogeneous medium (*κ* = 0.1 S/m globally, which is the average of white and grey matter conductivity). After the simulations, the Simbio/FieldTrip field is adjusted on the standard grid 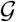 via interpolating function. As FastField relies on homogeneous media, conductivity value of 0.1 S/m is used in all simulations for *E*_1_.

Next, the divergence between *E*_1_ and *E*_2_ (Eq. 4) and the accuracy (Eq. 5) are calculated. Table 2 reports the accuracy values Acc(*E*_1_|*E*_2_). When considering FEM homogeneous condition, Acc(*E*_1_|*E*_2_) ∈ [0.9220; 0.9847] with an average value of 0.96. For FEM heterogeneous domain, Acc(*E*_1_|*E*_2_) ∈ [0.8038; 0.8582] with an average value of 0.83.

**Table 2.** Comparison of FastField with Simbio/FieldTrip e-fields. “Accuracy 1” refers to the homogeneous condition with *κ* = 0.1 S/m for all tissue types; “Accuracy 2” refers to the non-homogeneous condition, where conductivity values of 0.132 S/m and 0.08 S/m are used for grey and white matter respectively. In both cases a conductivity value of 0.1 S/m is applied in FastField. Amplitude values are in mA. Configuration values represent the percentage assigned to each contact of the electrode (contact sequences are numbered as in Fig. 5).

| Case | Accuracy 1 | Accuracy 2 | Electrode type | Amp | Configuration |
| --- | --- | --- | --- | --- | --- |
| 1 | 0.9220 | 0.8371 | Boston scientific vercise directed | 2.4 | 50,50,0,0,0,0,0,0 |
| 2 | 0.9278 | 0.8455 | Boston scientific vercise directed | 3.1 | 0,25,0,25,25,0,25,0 |
| 3 | 0.9819 | 0.8582 | Medtronic 3389 | 1.4 | 100,0,0,0 |
| 4 | 0.9623 | 0.8561 | Medtronic 3389 | 2.7 | 60,40,0,0 |
| 5 | 0.9605 | 0.8038 | Medtronic 3387 | 2.2 | 0,55,45,0 |
| 6 | 0.9847 | 0.8225 | Medtronic 3387 | 0.7 | 0,0,0,100 |
| 7 | 0.9636 | 0.8153 | Abbott/St Jude Medical Infinity Directed 6172 | 2.6 | 0,32,0,0,68,0,0,0 |
| 8 | 0.9523 | 0.8266 | Abbott/St Jude Medical Infinity Directed 6172 | 3.4 | 0,0,0,0,0,25,25,50 |

Finally, the Dice scores DS(VT_1_, VT_2_) are computed from Eq. 6 and are presented in Table 3. For the homogeneous condition, DS ∈ [0.9286; 0.9820] with an average value of 0.96. For non-homogeneous condition, DS ∈ [0.8667; 0.9335] with an average value of 0.92. Figure 8 shows several examples of VTA comparison, for different electrodes and contact configurations. FastField-based VTA isocontour is plotted in red, the Simbio/FieldTrip-based one is in blue.

**Table 3.** Dice score similarity of the FastField VTA with Simbio/FieldTrip VTA. “Dice score 1” refers to the homogeneous condition with *κ* = 0.1 S/m for all tissue types; “Dice score 2” refers to the non-homogeneous condition, where conductivity values of 0.132 S/m and 0.08 S/m for grey and white matter are used. In both cases, the conductivity values of 0.1 S/m is used in FastField. Amplitude values are in mA. Configuration values represent the percentage assigned to each contact of the electrode (electrodes are numbered as in Fig. 5).

| Case | Dice score 1 | Dice score 2 | Electrode type | Amp | Configuration |
| --- | --- | --- | --- | --- | --- |
| 1 | 0.9622 | 0.9393 | Boston scientific vercise directed | 2.4 | 50,50,0,0,0,0,0,0 |
| 2 | 0.9559 | 0.9349 | Boston scientific vercise directed | 3.1 | 0,25,0,25,25,0,25,0 |
| 3 | 0.9797 | 0.9529 | Medtronic 3389 | 1.4 | 100,0,0,0 |
| 4 | 0.9684 | 0.9190 | Medtronic 3389 | 2.7 | 60,40,0,0 |
| 5 | 0.9335 | 0.9468 | Medtronic 3387 | 2.2 | 0,55,45,0 |
| 6 | 0.9820 | 0.8667 | Medtronic 3387 | 0.7 | 0,0,0,100 |
| 7 | 0.9634 | 0.8735 | Abbott/St Jude Medical Infinity Directed 6172 | 2.6 | 0,32,0,0,68,0,0,0 |
| 8 | 0.9667 | 0.9165 | Abbott/St Jude Medical Infinity Directed 6172 | 3.4 | 0,0,0,0,0,25,25,50 |

**Fig. 8.**
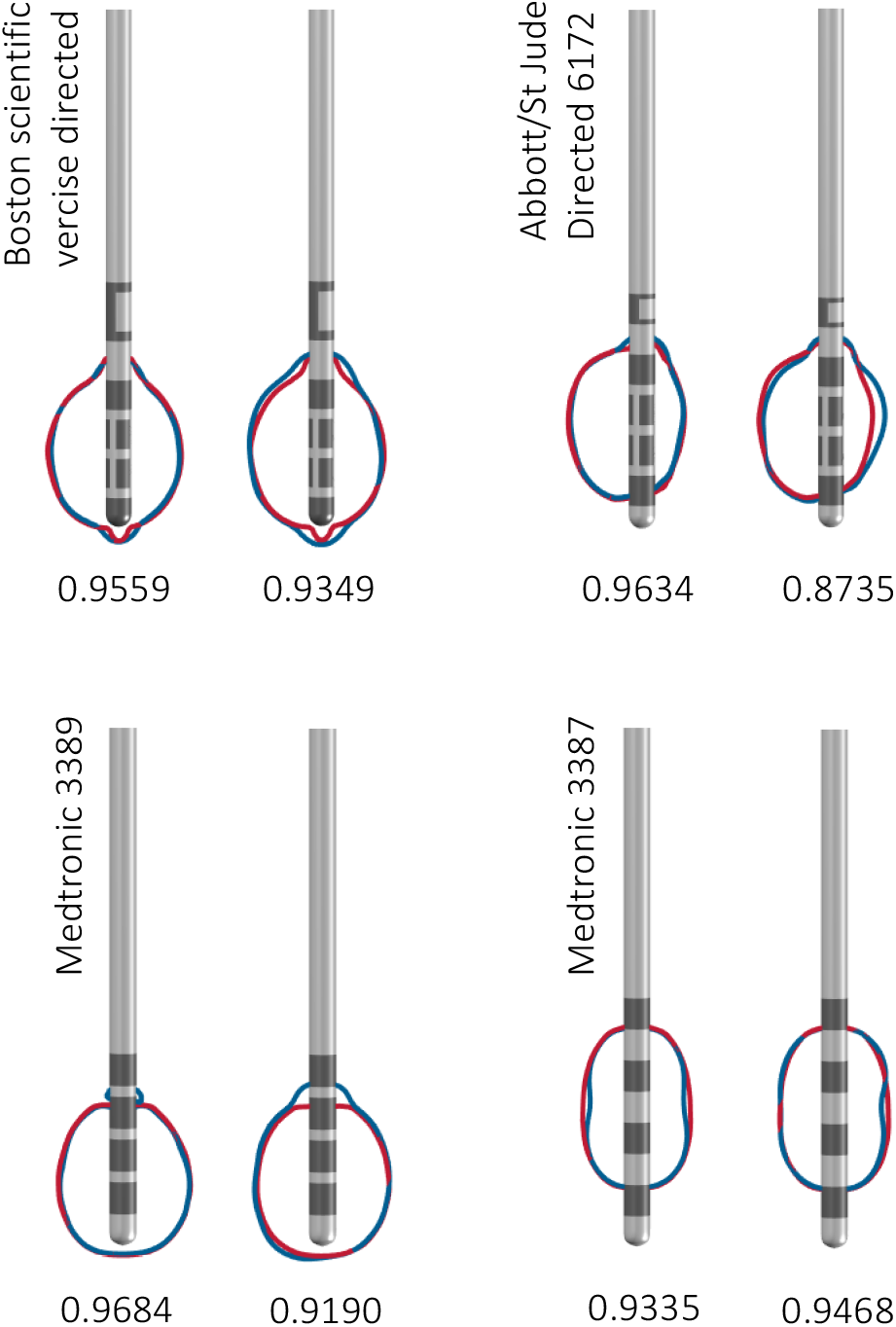
Comparison of FastField with Simbio/FieldTrip finite element model. Some example studies from Table 2 chosen for visualization (here, case 2, 4, 5, and 7). The FastField-based VTA is in red and the VTA simulated with Simbio/FieldTrip is in blue. For each electrode type, a couple of comparisons are shown: on the left, in a homogeneous domain (*κ* = 0.1 S/m for both FastField and Simbio/FieldTrip simulations); on the right, heterogeneous domain for Simbio/FieldTrip (*κ* = 0.08 S/m and 0.132 S/m for white and grey matter) and homogeneous *κ* = 0.1 S/m for FastField simulations. The Dice scores of the two VTA comparison is written under each figure. In this figure, the iso-surface of 0.2 V/mm (VTA) is shown as the VTA.

### 3.3. Case study 1

We consider a Parkinson patient with the STN target area. The electrode used is Boston scientific vercise directed; it is not placed inside, rather right next to the target. FastField is used to tune the parameters to direct the VTA towards the STN area. Rapid response from the algorithm allows to test different parameter configurations efficiently (in ~ 0.2 s). As a result, the tuned stimulation amplitude is 1.8 mA and the weighted configuration to deliver the energy is: 20% on Contact 1 and 80% on Contact 2 of the electrode. Fig. 9a reports the VTA obtained from the tuned e-field and the target region. An electric field with the tuned settings is simulated with Lead-DBS Simbio/FieldTrip (on non homogeneous medium) and compared to the result from FastField. Their relative accuracy (Eq. 5) equals Acc(*E*_1_|*E*_2_) = 0.8301. Fig. 9b shows a direct comparison of VTA isocontours (blue and red color, respectively). The Dice score for the VTA comparison (Eq. 6) is DS(VT_1_, VT_2_) = 0.9277.

**Fig. 9.**
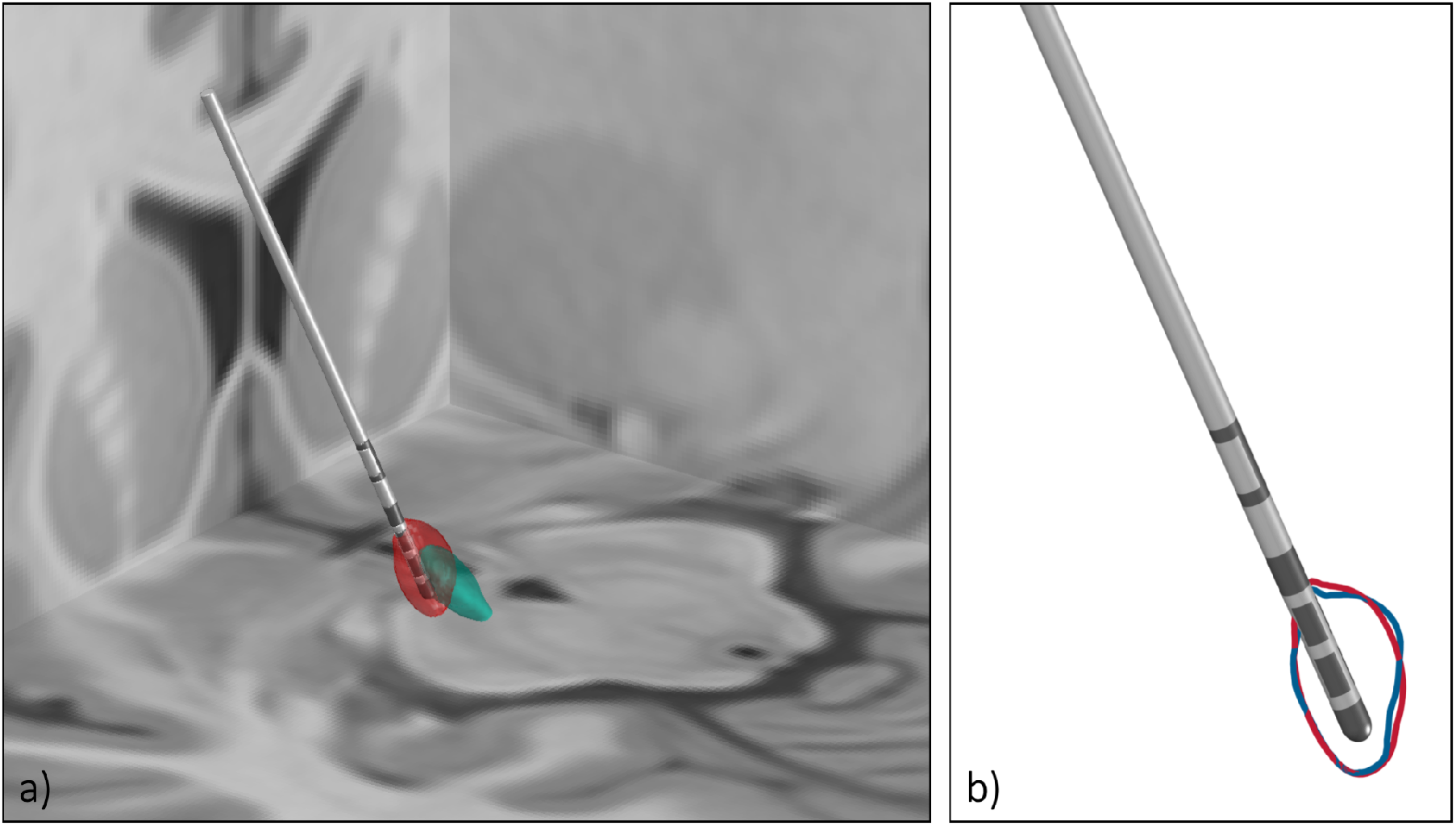
Clinical case study 1. A Parkinson patient with target structure STN. **a)** The approximated field with FastField. 20% of the energy comes from contact 1 and 80% from contact 2. Input amplitude is 1.8 mA. The e-field is in red and the STN is in green. **b)** Comparison of FastField with Simbio/FieldTrip for the same setting as in part (a). The e-field approximated with FastField is in red and the e-field simulated by Simbio/FieldTrip is in blue. The accuracy between the two fields is 0.8301. The Dice score for the two VTA is 0.9277.

### 3.4. Case study 2

Here, we consider a Post-Traumatic Tremor patient with internal globus pallidus (GPi) as target area. Medtronic 3389 electrode is used. The electrode was localized close to GPi. As in Case study 1, different setting configurations are tested efficiently using FastField to find an optimum. Eventually, Contact 4 (*w* = 100%, *A* = 2.5 mA) is identified as the appropriate setting for effective stimulation of GPi, while avoiding GPe to minimize possible side effects (Baizabal-Carvallo and Jankovic, 2016). Comparing Fastfield with Simbio/FieldTrip (non homogeneous domain) results in a relative accuracy of Acc(*E*_1_|*E*_2_) = 0.8686. Figure 10a represents the estimated output, i.e. the tuned e-field next to the target region. Figure 10b compares VTA results from FastField (red) and Simbio/FielTrip (blue) on the same tuned settings. In this case, DS(VT_1_, VT_2_) = 0.9200.

**Fig. 10.**
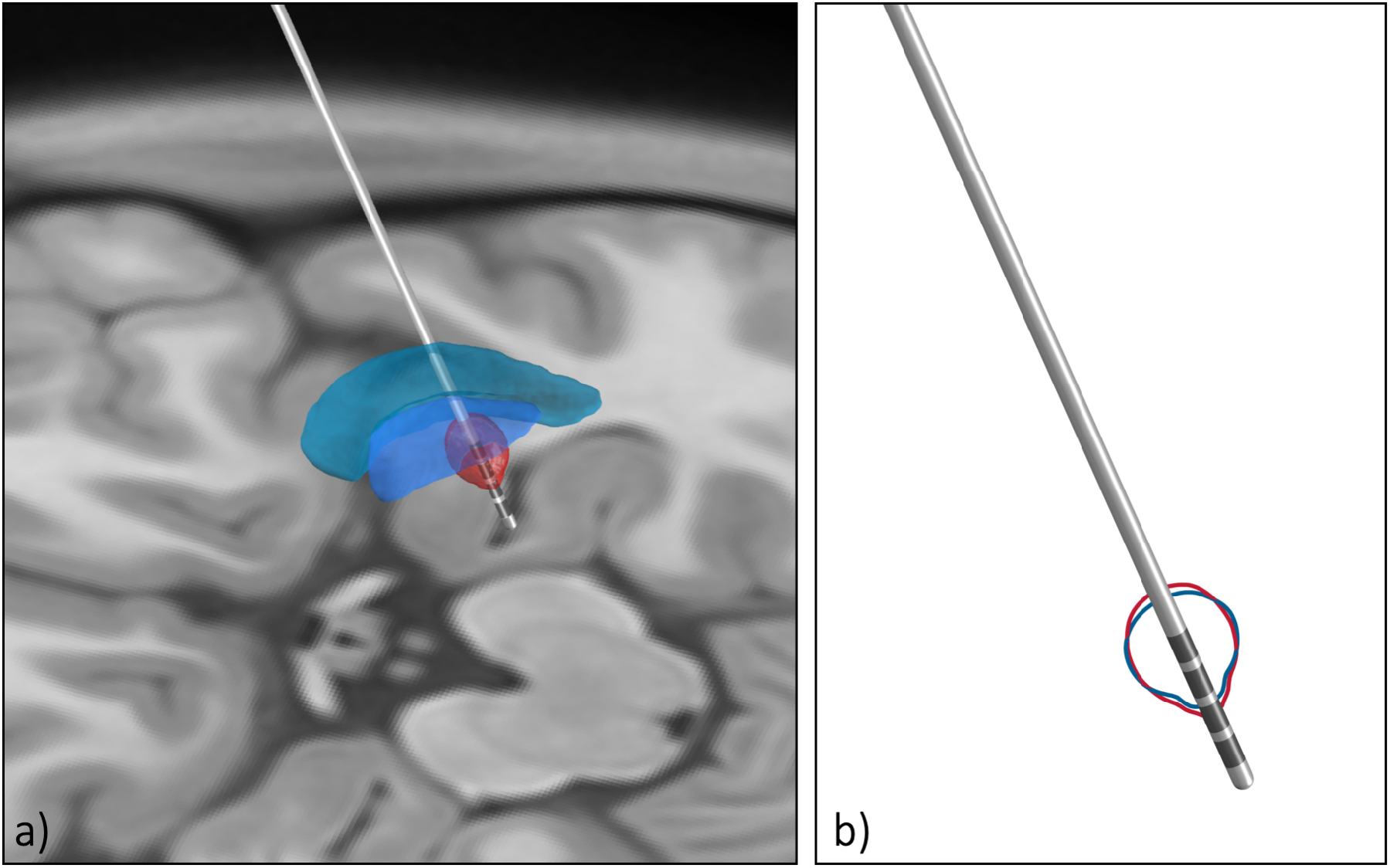
Clinical case study 2. A post-Traumatic Tremor patient with target structure GPi. a) The approximated field with FastField. 100% of the energy on contact 4 with the amplitude of 2.5 mA. The e-field is red and the GPi is blue, and GPe in green b) The comparison of FastField with Simbio/Field trip for the same setting as part a. The e-field approximated with FastField is in red and the e-field simulated by Simbio/FieldTrip is in blue. The similarity between the two field is 0.8686. The Dice score for the VTA comparison is 0.9200.

### 3.5. Case study 3

To show the use of DBS for psychiatric diseases, we also consider an Anorexia nervosa patient. In this case, nucleus accumbens (NAc) is identified as the target of interest. The electrode is Boston scientific vercise. As in previous case studies, different setting configurations are tested efficiently using FastField to find an optimal coverage of the NAc. Eventually, Contacts 2 (*w* = 15%), 3 (*w* = 75%), and 4 (*w* = 10%) are chosen with input current *A* = 2.2 mA. Comparison of Fastfield with Simbio/FieldTrip (non homogeneous) results in a relative accuracy of Acc(*E*_1_|*E*_2_) = 0.8603. Figure 11a shows the estimated tuned e-field nearby the target region. Figure 11b compares VTA results from FastField (red) and Simbio/FielTrip (blue) on the same tuned settings. In this case, DS(VT_1_, VT_2_) = 0.9302.

**Fig. 11.**
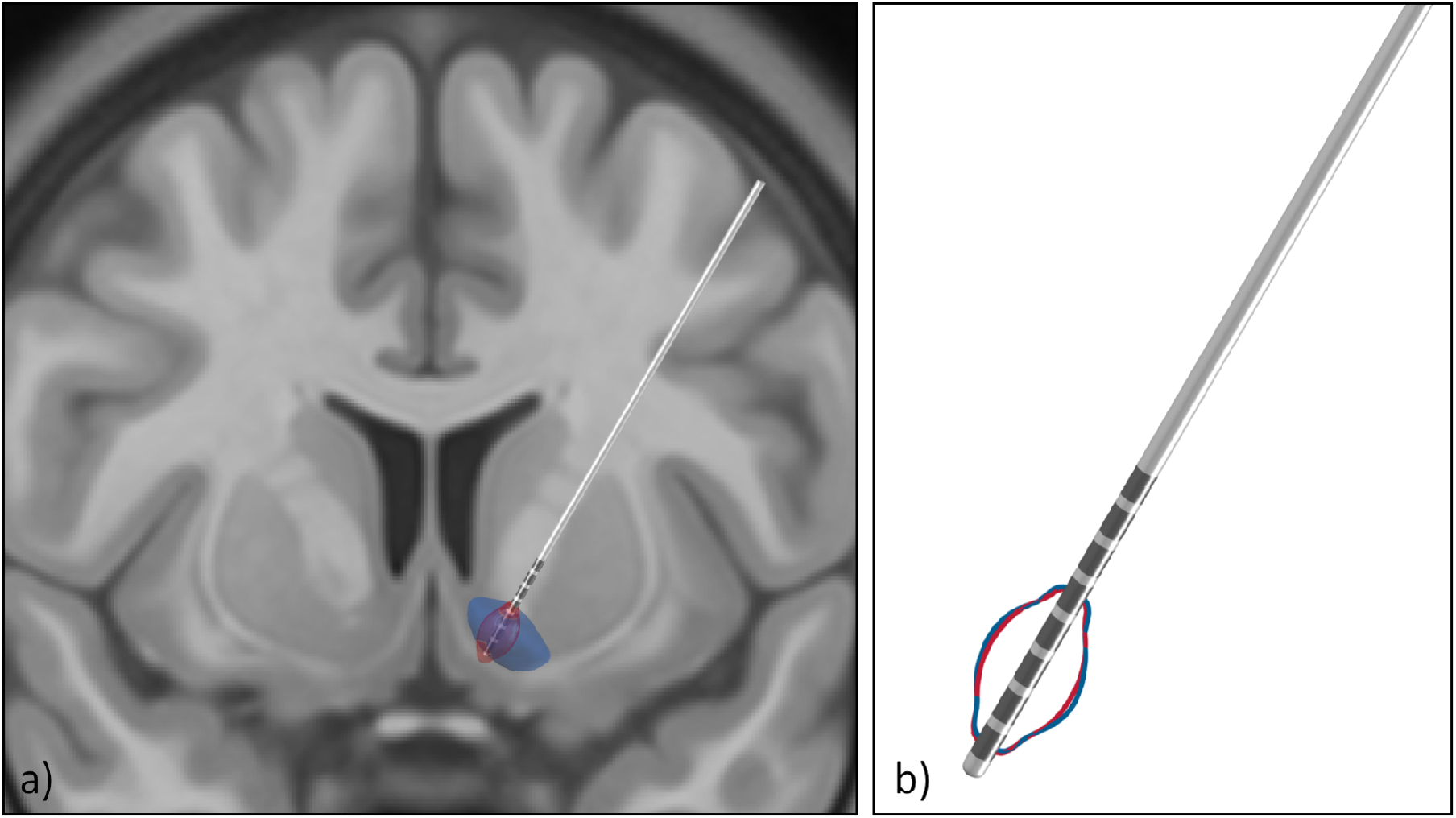
Clinical case study 3. An Anorexia nervosa patient with target structure nucleus accumbens. a) The approximated field with FastField. 10% of the energy on contact 2, 75% on contact 3, and 15% on contact 4, with the amplitude of 2.2 mA. The e-field is red and the nucleus accumbens is green, putamen in green, and caudate in purple. b) The comparison of FastField with Simbio/Field trip for the same setting as part a. The e-field approximated with FastField is in red and the e-field simulated by Simbio/FieldTrip is in blue. The similarity between the two field is 0.8603. The Dice score for the VTA comparison is 0.9302.

## 4. Discussion

We have introduced a toolbox to simulate the DBS electric field for a variety of electrode types. The toolbox was validated by comparing the results with a FEM model in a template space and clinical case studies.

### 4.1. Accuracy

To interpret the error index appropriately (Eq. 4), we contrast it with the measures uncertainty. This is due to real device resolution on input parameters. For instance, resolution of most of DBS devices is *σ_A_* =0.1 mA for the input amplitude value *A* (e.g. from Medtronic manual^1^). This is necessarily propagated by the algorithms. The corresponding uncertainty *σ_E_* on estimated e-field *E* is calculated for each case study by considering (*A*±*σ_A_*) for FEM-based models. Likewise, we evaluate Dice score (*DS_σ_*) for the two volumes computed from (*E* + *σ_E_*) and (*E* − *σ_E_*). For Case study 1, *A* = (1.8 ± 0.1) mA. The uncertainty associated to the output e-field is *σ_E_* = ±0.1103 V/mm. This is a realistic benchmark to contrast Err(*E*_1_|*E*_2_) with. In this case, we recall that Err(*E*_1_|*E*_2_) = 0.1699. Furthermore, we evaluate the Dice score on uncertainty VTAs, that equals DS_*σ*_ = 0.9114. This is even lower than DS(VT_1_, VT_2_) = 0.9277 as in Sec. 3.3.

Results for Case study 2 and 3 are consistent. For Case 2, Err(*E*_1_|*E*_2_) = 0.1314 while *σ_E_* = 0.0833; DS(VT_1_, VT_2_) = 0.9200 while DS_*σ*_ = 0.9322. For Case 3, Err(*E*_1_|*E*_2_) = 0.1397 and *σ_E_* = 0.0952; DS(VT_1_, VT_2_) = 0.9302 and DS_*σ*_ = 0.9048.

Hence, by recalling that other physical uncertainties (e.g. over pulse width and frequency) may further propagate the device uncertainty, we confidently conclude that, despite its approximation, FastField may serve as a reliable model for practical use.

### 4.2. Time efficiency

In terms of the computational time, Fastfield is more efficient than any finite element model. In fact, the algorithmic complexity of FastField is *O*(*N*), while that of a FEM is *O*(*N^a^*) where *a* usually varies between 2 and 3 (Liu and Quek, 2013). Consequently, as *N* ∝ *dim*^3^, FastField would scale as *O*(*N*^3^) and FEM as *O*(*N*^6^) (at best) when doubling the grid precision on every direction.

As a proof of concept, we estimate the CPU-time necessary to complete a simulation with FastField and with Simbio/FieldTrip. We use the same laptop for both (Macbook Pro, 2.3GHz Intel Core i5, 16 GB memory). For Simbio/FieldTrip, the whole computation (from stating the inputs to getting the VTA output) takes on average 400 seconds. Setting the meshed domain and assigning conductivity values is particularly demanding, as it accounts for about 65% of the whole procedure. Without considering this first step, the average computation time is about 140 s.

On the other hand, FastField avoids the expensive preliminary steps as it relies on the standard library to set the domain. Overall, simulating electric field and VTA takes about 0.2 seconds, 3 orders of magnitude less than with a FEM.

Augmented time performance in estimating the electric field is beneficial for many applications. For instance, in an optimization problem to tune the initial settings according to the target region. In such problem, the e-field is evaluated multiple times to test different settings towards the optimum. Without even considering the generation of the meshed domain, FastField saves around 140 seconds in each iteration, resulting in almost 4 hours after 100 iterations.

Another example where FastField is possibly beneficial is during clinical practice, for each time the physician changes the DBS parameter and evaluates the effect of new settings on neural tissue. In this case, enhanced computational speed could improve the user’s experience.

### 4.3. VTA model

The VTA activation model can be potentially used as a standalone function for direct use in any VTA simulation. However, caution is recommended when changing input voltage, as the original data for the fitting was taken at 3*V* (Åström et al., 2015). We conjecture the model to be extendable to other values, given that its functional dependence does not include input voltage explicitly. Further studies are suggested on this aspect.

For convenient use and to fosters reproducible research, open source Matlab functions of the model are provided.

### 4.4. Limitations

Given the main advantages of FastField, we acknowledge its main limitation, that the simulated domain is treated as a homogeneous medium. Despite such approximation being essential to diminish the computational burden and thus boosting the speed, considering different conductivity values for different brain tissues would eventually increase the precision of the method. Moreover, we notice that there exists a big difference among the conductivity values used in recent DBS field simulation studies (cf. Table 1), which is also discussed in (McCann et al., 2019). This is supposedly due to relevant difference between the conductivity values of different patients (Koessler et al., 2017). Therefore, the conductivity value is a free parameter in FastField, to be tuned by the user. We hope that further studies will improve the estimation of the patient’s specific conductivity values and that future work will enable better models and turn the homogeneous approximation superfluous soon.

We finally remark that not all the existing electrode types are currently supported in the current FastField release: twelve electrode types from four different vendors are now considered. Others can be easily added in future, as FastField allows easy embedding of different geometries.

## 5. Conclusion

FastField is a user-friendly toolbox to approximate the DBS electric field in a fast and accurate way. The precision of the method is comparable to that of a FEM model with the assumption of a homogeneous medium in the vicinity of the electrode, which is often sufficient for practical use. Its time performance is ~ 1000 times faster than a FEM model, which makes it useful for many applications in abstract studies and clinical practice. FastField considers the most relevant parameters for the stimulation, enriching their set with pulse width and axon diameter for VTA approximation (usually neglected in other studies). Hence, we hope it will foster insightful and reproducible studies on the effect of DBS stimulation on brain networks.

## Code availability

FastField Matlab code and graphical user interface are available under GNU licence on https://github.com/luxneuroimage/FastField.

VTA heuristic model as standalone function is available on https://github.com/luxneuroimage/ApproXON. An integration of FastField to the LeadDBS deep brain stiumlation toolbox is going to be provided at (https://github.com/netstim/leaddbs).

## Data Disclosure

Anonymized data for the case studies were obtained from Centre Hospitalier du Luxembourg following Ethics approval CNER 201804/06 (EINSDBS)

## Acknowlegement

Authors would like to thank the Lead-DBS team for providing the Lead-DBS software and related resources.

## CRediT authorship contribution statement

**Mehri Baniasadi:** Conceptualization, Methodology, Software, Validation, Formal analysis, Investigation, Writing - Original Draft, Writing - Review & Editing, Visualization. **Daniele Proverbio:** Methodology, Software, Formal analysis, Writing - Original Draft, Writing - Review & Editing, Visualization. **Jorge Gonçalves:** Writing - Review & Editing, Supervision, Project administration, Funding acquisition. **Frank Hertel:** Supervision, Project administration, Writing - Review & Editing. **Andreas Husch:** Conceptualization, Methodology, Software, Formal Analysis, Investigation, Writing - Review & Editing, Supervision, Project administration.

## Footnotes

1 http://www.neuromodulation.ch/sites/default/files/pictures/activa_PC_DBS_implant_manuel.pdf

